# Pangenome databases provide superior host removal and mycobacteria classification from clinical metagenomic data

**DOI:** 10.1101/2023.09.18.558339

**Authors:** Michael B. Hall, Lachlan J.M. Coin

**Affiliations:** Department of Microbiology and Immunology, Peter Doherty Institute for Infection and Immunity, The University of Melbourne, Melbourne, Australia

**Keywords:** host removal, metagenomics, *Mycobacterium tuberculosis*, taxonomic classification, benchmark

## Abstract

**Background:** Culture-free real-time sequencing of clinical metagenomic samples promises both rapid pathogen detection and antimicrobial resistance profiling. However, this approach introduces the risk of patient DNA leakage. To mitigate this risk, we need near-comprehensive removal of human DNA sequence at the point of sequencing, typically involving use of resource-constrained devices. Existing benchmarks have largely focused on use of standardised databases and largely ignored the computational requirements of depletion pipelines as well as the impact of human genome diversity.

**Results:** We benchmarked host removal pipelines on simulated Illumina and Nanopore metagenomic samples. We found that construction of a custom kraken database containing diverse human genomes results in the best balance of accuracy and computational resource usage. In addition, we benchmarked pipelines using kraken and minimap2 for taxonomic classification of *Mycobacterium* reads using standard and custom databases. With a database representative of the *Mycobacterium* genus, both tools obtained near-perfect precision and recall for classification of *Mycobacterium tuberculosis*. Computational efficiency of these custom databases was again superior to most standard approaches, allowing them to be executed on a laptop device.

**Conclusions:** Nanopore sequencing and a custom kraken human database with a diversity of genomes leads to superior host read removal from simulated metagenomic samples while being executable on a laptop. In addition, constructing a taxon-specific database provides excellent taxonomic read assignment while keeping runtime and memory low. We make all customised databases and pipelines freely available.

## Introduction

*Mycobacterium tuberculosis* is the bacterium that causes tuberculosis, which is a leading cause of death globally[1]. Tuberculosis is an ancient airborne disease that predominantly affects the lungs [2]. Whole-genome sequencing (WGS) with Illumina and Oxford Nanopore Technologies platforms is increasingly being used for *M. tuberculosis* diagnostic applications such as drug resistance prediction, lineage determination, and identification of putative transmission clusters[3, 4, 5, 6]. Currently, most WGS applications for *M. tuberculosis* rely on culturing of the bacterium on either solid or liquid media-which can take days to weeks. Sequencing of *M. tuberculosis* directly from patient sample (sputum) is a much desired solution as it provides faster time to results and does not require the infrastructure necessary for culture.

A number of issues exist when sequencing *M. tuberculosis* direct from sputum - and indeed, any human-associated metagenomic sample. *M. tuberculosis* genomic DNA is generally scarce in such metagenomic samples, with host and other bacterial DNA dominating[7, 8]. Capture of *M. tuberculosis* DNA can be attempted during sample preparation, or computationally during sample quality control. We focus here on the computational extraction of *M. tuberculosis* DNA from metagenomic sputum samples. Previous work has shown that removal of non-*M. tuberculosis* sequencing reads from metagenomic samples is crucial for reducing false-positive and -negative variant calls in downstream analyses[9], even for samples with low levels of contamination. One solution that is common in bioinformatic pipelines is contamination removal using either alignment tools or taxonomic classifiers. Kraken[10] is a popular taxonomic classifier used for this purpose, with standardised databases generally being used [9, 11]. Alignment to the *M. tuberculosis* H37Rv reference genome is another favoured choice [12, 13], although this approach has been shown to still propagate false variant calls[9]. A more robust alignment approach is competitive mapping, where alignment databases are constructed to include common contaminants[14]. The idea with competitive mapping is that including a variety of species decreases the likelihood that reads from organisms with similar sequence will incorrectly map to your organism of interest.

While *M. tuberculosis* contamination removal and read classification have been assessed previously[9, 15], these studies have focused on Illumina sequencing data. However, Nanopore data is becoming a popular choice for *M. tuberculosis* metagenomic studies[8, 16]. As Nanopore sequencing can be performed on a laptop, analysis pipelines should use computational resources that make them executable on such machines. Goig *et al*. recommended use of the standard kraken database for removal of contamination and extrication of *M. tuberculosis* DNA[9], however, this database uses upwards of 70GB of memory; an amount which is not available on laptops, or indeed most desktop computers, thus requiring users to have access to high performance computing resources.

In this study, we assess a variety of tools for the removal of human reads in metagenomic samples, as well as the classification of *M. tuberculosis* reads, from both simulated Illumina and Nanopore sequencing platforms. We note that the removal of host reads is applicable to any human-associated metagenomic samples. We place a strong emphasis on computational resource usage in addition to accuracy. While we assess standard databases, we also create custom databases for both tasks and make them freely available with this work. This curation of custom databases allows us to keep computational resource usage low, while maintaining high read classification accuracy (higher than existing methods in most cases).

## Results

We generated an *in silico* metagenomic readset from a variety of organisms (human, bacteria, virus) at ratios commonly seen in *M. tuberculosis* metagenomic samples (see Generating *in silico* metagenomic reads)[8]. After removing Nanopore reads shorter than 500bp or with an ambiguous base, we were left with 234984 reads with a total of 2.48 gigabases[17]. For Illumina, after removing reads with an ambiguous base, we retained 2753282 read pairs with a combined total of 826 megabases[18].

### Removal of human reads

A primary focus of this work is capturing *M. tuberculosis* complex reads from metagenomic samples, with a secondary aim of doing this in low-resource computational settings (e.g. on a laptop). In general, for these types of samples, human reads are undesirable and so their removal is an important first step. While human read removal could be built into the broader classification of such a sample, removal *ab initio* reduces memory requirements (smaller, modular databases can be used), decreases file sizes, and therefore runtimes in downstream analyses, and avoids accidental patient DNA ending up in data submitted to public archives. In addition, host read removal is a task common to most metagenomic applications, so the separate assessment is likely to be of interest to a wider audience.

We benchmarked six configurations from both *k*-merand alignment-based approaches for classifying reads as being human or not (see Human read removal). The *k*-mer-based methods include the human read removal tool (HRRT)[19] and kraken[10] with the default human database[20] and a database we built from the 97 assemblies used by the Human Pangenome Reference Consortium (HPRC)[21, 22]. The alignment-based approaches include minimap2[23], Hostile[24] (which uses minimap2 and does some extra filtering), and minimap2 followed by winnowmap[25] (*miniwinnow*; note, this option was not applicable for Illumina data). The reference used for each alignment method was the CHM13v2 assembly[26] (NCBI RefSeq accession GCF_009914755.1) plus human leukocyte antigen (HLA) sequences[24]. Note, the human reads were simulated from the Korean reference genome (KOREF_S1v2.1 accession GCA_020497085.1)[27] which is not present in any of the human removal databases used here.

#### Nanopore

Table 1 presents the computational and accuracy performance for each method on Nanopore data. From this, we see that all methods have very high recall and precision. Hostile and miniwinnow produce the best balance of precision and recall, with an F-score of 0.99991. Kraken (both databases) was an order of magnitude faster than HRRT, Hostile and miniwinnow, and at least six times faster than minimap2. Lastly, the *k*-mer-based classification methods had much lower peak memory usage than their alignment-based competitors, with memory usage low enough to be suitable for operation on most laptop devices (< 8GB).

**Table 1.**
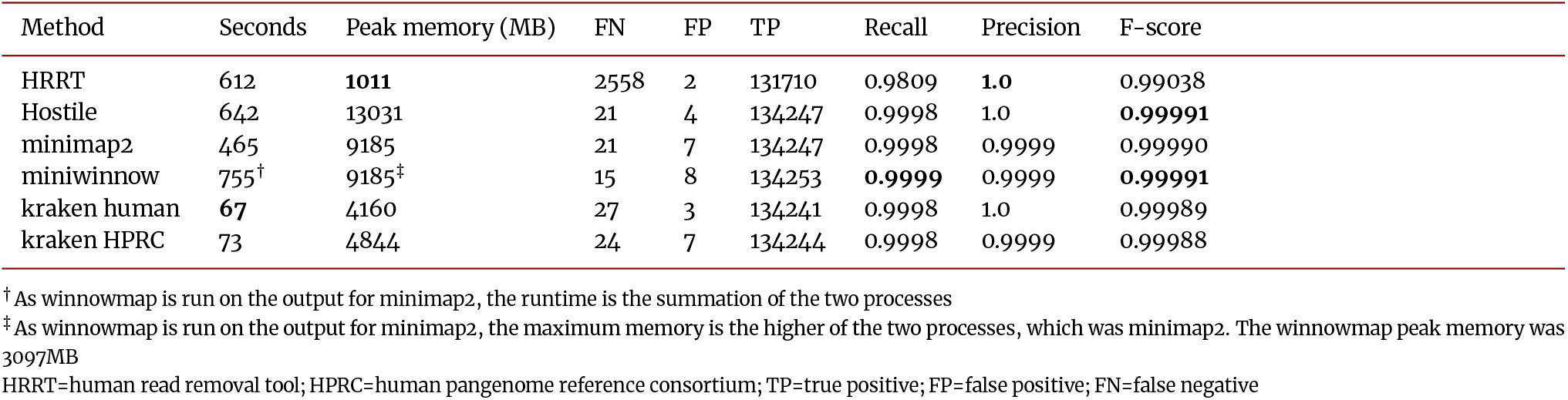
Performance of human read classification - Nanopore.

Importantly, kraken did not classify any *Mycobacterium* reads as being human. However, all other methods did, albeit a very small number (Table 2).

**Table 2.** *Mycobacterium* false positive counts during human read classification - Nanopore.

|  | *M. tuberculosis* | *M. kansasii* | *M. ulcerans* |
| --- | --- | --- | --- |
| HRRT | 0 | 0 | 2 |
| Hostile | 1 | 2 | 0 |
| minimap2 | 1 | 3 | 0 |
| miniwinnow | 2 | 3 | 0 |
| kraken human | 0 | 0 | 0 |
| kraken HPRC | 0 | 0 | 0 |

#### Illumina

Table 3 presents the results for human read removal on Illumina data. As expected, the Illumina results have lower recall than Nanopore, with minimap2 providing the highest recall (0.9837). Despite the lower recall, all methods performed exceptionally for precision, with HRRT producing no false positives (FPs), and minimap2 producing the highest F-score (0.9918). The runtime of the methods on Illumina data is much the same as for Nanopore data, with kraken being an order of magnitude faster than the other methods. Aside from minimap2, peak memory usage for all methods is low enough to be suitable for operation on most laptop devices (< 8GB).

**Table 3.** Performance of human read classification - Illumina.

| Method | Seconds | Peak memory (MB) | FN | FP | TP | Recall | Precision | F-score |
| --- | --- | --- | --- | --- | --- | --- | --- | --- |
| HRRT | 313 | 1042 | 268479 | 0 | 2462217 | 0.9017 | 1.0 | 0.9483 |
| Hostile | 234 | 3781 | 51966 | 4 | 2678730 | 0.9810 | 1.0 | 0.9904 |
| minimap2 | 383 | 11994 | 44544 | 137 | 2686152 | 0.9837 | 0.9999 | 0.9918 |
| kraken human | 20 | 4228 | 71684 | 20 | 2659012 | 0.9737 | 1.0 | 0.9867 |
| kraken HPRC | 20 | 4927 | 58196 | 34 | 2672500 | 0.9787 | 1.0 | 0.9892 |

HRRT=human read removal tool; HPRC=human pangenome reference consortium; TP=true positive; FP=false positive; FN=false negative

Minimap2 was the only method that classified *Mycobacterium* Illumina reads as being human: 11 *M. tuberculosis*, 1 *M. ulcerans*, and 1 *M. kansasii*. In addition to those 13 *Mycobacterium* reads, the most common genera erroneously classified as human by minimap2 were *Xanthomonas* (*n* = 7) and *Streptococcus, Clostridium, Clostridioides*, and *Acinetobacter* (all *n* = 5). All four of the Hostile FPs were from Human endogenous retrovirus K (HERV-K). Four of the FPs from both kraken databases were also HERV-K, along with six Epstein-Barr virus reads and four *Campylobacter* against the HPRC database.

Almost all of the missed human reads (false negatives (FNs)) for all methods were from unplaced scaffolds in the KOREF_S1v2.1 genome.

### Classification of *Mycobacterium* reads

Having assessed the removal of human reads from *M. tuberculosis* metagenomic samples, we turn to read classification. In particular, we focus on being able to classify *M. tuberculosis* reads to the correct species, with as few false-positives and -negatives as possible. For this section of the analysis, we use reads that were unclassified by kraken on the HPRC database (i.e., non-human reads). In total, we are left with 100733 Nanopore reads (460 megabases) and 1417015 Illumina read pairs (424 megabases).

We benchmarked kraken and minimap2 for read classification, using three different databases for each tool. For kraken, we used the standard database (*standard*) and a version of the standard database that is capped at 8GB (*standard-8*). In addition, we created a *Mycobacterium*-specific database which contains a RefSeq genome from each species in the *Mycobacteriaceae* family plus a variety of other genera (see *Mycobacterium* read classification)[28]. For minimap2 we created an *M. tuberculosis*-specific database (*MTB*) using the *M. tuberculosis* H37Rv reference genome and 17 high-quality *M. tuberculosis* genomes from lineages 1-6 [29, 30]. We also created a *Mycobacterium*-specific database (*Mycobacterium*) with a RefSeq genome from each species in the *Mycobacterium* genus[31]. And lastly, we used the decontamination database from the Clockwork pipeline [14], but also added the 17 high-quality *M. tuberculosis* genomes[32].

#### Nanopore

Table 4 shows the *M. tuberculosis* read classification results for Nanopore data. From this we see that minimap2 with the Clock-work and *Mycobacterium* databases had the highest recall with zero and one false negative call, respectively. Kraken (all databases) and minimap2 with the *Mycobacterium* database have the highest precision (1.0). Minimap2 with the *Mycobacterium* database gave the highest F-score (balance between recall and precision) of 1.0 producing only one FN and three FPs. Kraken with the standard-8 database was the fastest method, however all methods ran in less than four and a half minutes. Memory consumption varied quite a bit across methods, with minimap2’s MTB database using the lowest at 2050MB. Notably, kraken standard would not be executable on a computer with 16GB of memory, whilst minimap2 Clockwork is very close to this limit.

**Table 4.** Performance of *M. tuberculosis* read classification - Nanopore.

| Method | Seconds | Peak memory (MB) | FN | FP | TP | Recall | Precision | F-score |
| --- | --- | --- | --- | --- | --- | --- | --- | --- |
| minimap2 Clockwork | 235 | 14220 | 0 | 48 | 60257 | 1.0 | 0.9992 | 0.9996 |
| minimap2 MTB | 196 | 2050 | 1786 | 772 | 58471 | 0.9704 | 0.987 | 0.9786 |
| minimap2 *Mycobacterium* | 251 | 7939 | 1 | 3 | 60256 | 1.0 | 1.0 | 1.0 |
| kraken standard | 83 | 68512 | 15620 | 0 | 44638 | 0.7408 | 1.0 | 0.8511 |
| kraken standard-8 | 22 | 7944 | 34973 | 0 | 25285 | 0.4196 | 1.0 | 0.5912 |
| kraken *Mycobacterium* | 29 | 8385 | 284 | 1 | 59974 | 0.9953 | 1.0 | 0.9976 |

#### Illumina

Table 5 presents the *M. tuberculosis* read classification results for Illumina data. The precision and recall best-performers are similar to the Nanopore results, with minimap2 Clockwork having the highest recall and kraken (all databases) the highest precision. Minimap2 Clockwork and *Mycobacterium* have the equal-highest F-score at 0.9997. In terms of runtime, kraken standard-8 was again the fastest, however all methods ran in under one and a half minutes. Memory consumption was again similar to Nanopore data, with minimap MTB using the lowest (804MB) and minimap2 Clockwork and kraken standard using more than 16GB.

**Table 5.** Performance of *M. tuberculosis* read classification - Illumina.

| Method | Seconds | Peak memory (MB) | FN | FP | TP | Recall | Precision | F-score |
| --- | --- | --- | --- | --- | --- | --- | --- | --- |
| minimap2 Clockwork | 87 | 21309 | 24 | 204 | 350944 | 0.9999 | 0.9994 | 0.9997 |
| minimap2 MTB | 29 | 804 | 17516 | 2194 | 315836 | 0.9475 | 0.9931 | 0.9697 |
| minimap2 *Mycobacterium* | 48 | 10402 | 148 | 58 | 384122 | 0.9996 | 0.9998 | 0.9997 |
| kraken standard | 60 | 68345 | 279074 | 4 | 54522 | 0.1634 | 0.9999 | 0.281 |
| kraken standard-8 | 14 | 7900 | 315058 | 0 | 18276 | 0.0548 | 1.0 | 0.104 |
| kraken *Mycobacterium* | 19 | 8428 | 8960 | 8 | 324636 | 0.9731 | 1.0 | 0.9864 |

Minimap2 with the *Mycobacterium* database made 148 FNs and 58 FPs. Of these, all FNs were aligned to non-*M. tuberculosis* species, with 50 being assigned to *M. angelicum* and 36 to *M. shinjukuense*. No genera were over-represented in the 58 FPs, however, there were two human reads in this set.

## Discussion

We have performed a thorough evaluation of both human and *M. tuberculosis* read classification from simulated metagenomic mixtures with both Nanopore and Illumina sequencing data. We have focused on human and *M. tuberculosis* classification in separate steps for two reasons. First, human reads are never required in typical *M. tuberculosis* analysis and are almost never uploaded to public sequence archives due to legal and ethical reasons. Second, removing human reads from the outset reduces computational time and memory usage in subsequent analyses and reduces the required bandwidth if data is to be uploaded to a public archive or cloud computing platform. As Nanopore sequencing can be performed on a laptop, the ability to perform contamination removal and read classification on such computers is an important consideration-especially with regards to memory consumption.

Most *M. tuberculosis* WGS work is done from culture and therefore does not contain much contamination by design. However, the eventual goal for the community is to move towards WGS directfrom-sputum analysis. Taking such samples directly from the patient, without a culturing step, will speed up time-to-results dramatically but does result in increased levels of contamination from the patient and other commensal species. Reducing the number of reads falsely classified as *M. tuberculosis* is important for down-stream analyses. In particular, false non-*M. tuberculosis* reads can cause false positives and negatives during variant calling [9, 11]. As variant calling underpins many vital *M. tuberculosis* applications, such as transmission cluster detection and drug resistance prediction[4], host removal and read classification accuracy is critical.

We have tested standardised host removal methods and created a novel, custom kraken database from a diverse range of human genomes produced as part of the Human Pangenome Reference Consortium (HPRC) [21] (see Data availability). We find that using this HPRC kraken database strikes the best balance between computational speed, memory consumption and reduced false classifications. While it was not the method with the highest F-score (though it was very close) we believe it will be the most robust to use in different human populations across both Nanopore and Illumina data. Although, we do note that Hostile is also another strong option for use on Illumina data. While this analysis was applied in the context of *M. tuberculosis* metagenomic samples, removal of human data is not limited to such cases and the kraken database can be used in other metagenomic situations.

When comparing human read removal between the two sequencing technologies, we found Nanopore data resulted in 2-3% higher recall across the different pipelines and databases tested. Therefore, if host removal is a crucial component of one’s desired analysis, Nanopore warrants serious consideration as a sequencing choice.

For *M. tuberculosis* read classification, we tested some standard databases for minimap2 and kraken and also constructed customised *M. tuberculosis* databases (see Data availability). There is no clear best performer across all situations, however, the *Mycobacterium* minimap2 and kraken database are our two recommendations. The choice of which of these databases to use will be up to individual users and their computational resource availability and whether reduction of false positives or negatives is of more importance.

Although the read classification and custom databases were targeted at *M. tuberculosis*, this work should act as a guide for how to design such taxon-specific databases for other species.

In conclusion, we found that construction of custom human and *M. tuberculosis* databases improves human read removal and *M. tuberculosis* read classification in simulated metagenomic samples. We make all custom databases freely available, along with usage examples (see Data availability).

## Methods

### Generating *in silico* metagenomic reads

We simulated metagenomic Nanopore and Illumina sequencing reads to a mixture ratio that approximates that found in patient sputa[8], albeit with a slightly higher mycobacterial component. In total, 4.5 and 0.9 gigabases were generated for Nanopore and Illumina, respectively, at proportions: 46% each for bacteria and human, 6% *M. tuberculosis* complex (MTBC), and 1% each for virus and non-tuberculous mycobacteria (NTM).

The reference genomes that reads were simulated from for these groups were gathered as follows. The references for the virus group were obtained using kraken’s (v2.1.2)[10] `--download-library` functionality. The viral library was downloaded on June 15 2023. The human genome from which the reads were simulated was the Korean reference KOREF_S1v2.1 (RefSeq accession GCA_020497085.1)[27], with contigs shorter than 10kbp removed. The bacterial references were obtained by first downloading the bacteria library through kraken, followed by a subsampling due to the size (166Gb) of the resulting FASTA file. We subsampled the file by first removing sequences with a length < 50kbp. We then extracted each sequence into its own FASTA file under a directory for the genus of the sequence-excluding the *Mycobacterium* genus[33, 34]. Genera were randomly subsampled to contain a maximum of 1000 assemblies. Each genus was then reduced to a representative subset using Assembly Dereplicator (commit 2dfcb14)[35] by keeping only 10% of the assemblies for each genus (`-f 0.1`). The NTM references selected were *M. abscessus* (accession GCF_017190695.1), *M. avium* (GCF_020735285.1), *M. kansasii* (GCA_014701265.1), *M. ulcerans* (GCF_000013925.1), *M. intracellulare* (GCF_016756075.1), *M. terrae* (GCF_010727125.1), and *M. fortuitum* (GCF_001307545.1). The MTBC reference is a lineage 1 assembly (GCF_932530395.1).

We used Badreads (v0.4.0)[36] to produce the simulated Nanopore reads for each group, specifying the number of bases in the appropriate proportions mentioned above. For all groups, we specified no junk or random reads and 0.5% chimeric reads. In addition, for the MTBC, virus, and NTM groups we used a non-default length option `--length 4000,3000` to produce reads with mean length 4000bp and a standard deviation of 3000. Defaults were used for all other options (the default error model is trained on real R10.4.1 Nanopore reads from 2023).

Illumina reads were simulated with ART (v2016.06.05)[37], using a similar approach as for Nanopore to generate reads for each group. We simulated paired reads from a MiSeq v3 system (`-ss MSv3`) with a read length of 150, a mean fragment length of 250 and fragment length standard deviation 10 (`-l 150 -m 250 -s 10`).

We filtered the simulated Nanopore reads to remove any read with a length < 500bp or an ambiguous nucleotide (non-ACGT) and removed simulated Illumina reads with any ambiguous base.

### Human read removal

We tested six configurations for removal of human reads from our simulated Nanopore metagenomic dataset: the human read removal tool (HRRT; v2.1.0)[19] with the `-x -r` options to remove human reads instead of masking them and write them to a separate file; Hostile (v0.0.3)[24] using the minimap2 aligner option; minimap2 (v2.26)[23] with the `-x map-ont` option; the fourth configuration was running winnowmap (v2.0)[25] on the non-human reads from the minimap2 configuration mentioned previously, using the same options as minimap2 (we label this configuration *miniwinnow*); the remaining two configurations were using kraken with a two different databases. One was built using only the default human library that comes with the `--download-library human` option. This library contains the CHM13v2[26] and GRCh38 references -we term this configuration *kraken human*. The second database consists of all 97 assemblies from the Human Pangenome Reference Consortium[21] we call this database *kraken HPRC*.

For the Illumina dataset we used five configurations -the same methods as above, minus the miniwinnowapproach as it onlyworks for long reads. Hostile uses Bowtie2[38] as the read aligner when operating on Illumina data. For Illumina we used the `-x sr` option in minimap2 instead and in kraken we additionally used the `--paired` option.

We assess the performance of these configurations using precision, recall, F-score (harmonic mean of precision and recall), peak memory usage, and runtime (seconds). All tools were run using four threads for consistency. In the context of this analysis we define a true positive (TP) as a read that originates from a human genome and was classified as being a human read, a false positive (FP) as a non-human read that was classified as a human read, and a false negative (FN) as a human read that was not classified as human.

### *Mycobacterium* read classification

Two classification tools, kraken and minimap2, were evaluated with three different databases each. For kraken, we used the standard database[39] which contains complete RefSeq genomes from bacterial, archaeal, and viral domains, the human genome, and a collection of known vectors, along with a version of this database that is capped at 8GB (*standard-8*)[40]. These two databases were downloaded from https://benlangmead.github.io/aws-indexes/k2 and were built on 06/05/2023. The third kraken database used was a *Mycobacterium*-specific one. To generate this database, we used `genome_updater` (v0.6.3)[41] to download one RefSeq genome from each species in the *Mycobacteriaceae* family, plus one RefSeq genome from each species in the following genera: *Klebsiella, Escherichia, Salmonella, Enterobacter, Streptococcus, Staphylococcus, Pseudomonas, Xanthomonas*, and *Bifidobacterium*. We ensured that the genomes used to simulate NTM and MTBC reads were not included in this database. We then built the kraken database from these genomes using `kraken2-build --build` with default options.

For minimap2, one database (*MTB*) contains the *M. tuberculosis* H37Rv reference genome (RefSeq accession GCF_000195955.2), plus 17 high-quality *M. tuberculosis* references from lineages 1-6 [29]. The second database (*clockwork*) contains common sputum contaminants along with a selection of NTM genomes and H37Rv -as used in the Clockwork pipeline[14]. In addition, we added the 17 high-quality *M. tuberculosis* genomes to this collection. The third minimap2 database we built was a *Mycobacterium*-specific one. For this database, we used `genome_updater` to download one RefSeq genome from each leaf node in the *Mycobacterium* genus[33, 34] plus the 17 high-quality *M. tuberculosis* assemblies.

The reads used for classification were those deemed non-human by kraken against the HPRC database (see Human read removal). We ran minimap2 with the present option `-x sr` for Illumina and `-x map-ont` for Nanopore along with base-level alignment and no secondary alignments (`-c --secondary=no`). We use default options for kraken classification, except for the `--paired` option for Illumina data.

We assess the performance of these tools and databases with the same metrics and threads as human read removal (Human read removal). In the context of this analysis we define a true positive (TP) as a read that originates from a *M. tuberculosis* genome and was classified as being a *M. tuberculosis* read, a false positive (FP) as a non-*M. tuberculosis* read that was classified as a *M. tuberculosis* read, and a false negative (FN) as a *M. tuberculosis* read that was not classified as *M. tuberculosis*.

## Availability of source code and requirements

All code to perform the analysis in this work and produce the custom databases can be found at the project described below. All steps in the pipeline were executed in reproducible remote containers or conda environments, which are listed in the project configuration file within the repository.

- Project name: Classification benchmark
- Project home page: classification_benchmark https://github.com/mbhall88/
- Operating system(s): Platform independent
- Programming language: Snakemake, Python, Perl, and Bash
- License: MIT

This repository also contains example usage of the custom databases and how they can be used to classify reads for *M. tuberculosis*.

## Data availability

The datasets supporting the results of this study are available in Zenodo, and have been cited in Results where first mentioned. They include the simulated Nanopore[17] and Illumina[18] reads, the human-only[20] and HPRC[22] kraken databases, the *Mycobacterium*-specific kraken database[28], plus the Clockwork[32], *M. tuberculosis*-specific[30], and *Mycobacterium*-specific databases[31] used by minimap2. The standard[39] and standard-8[40] kraken databases were downloaded from https://benlangmead.github.io/aws-indexes/k2.

## Declarations

## List of abbreviations

FN: false negative
FP: false positive
HERV-K: Human endogenous retrovirus K
HLA: human leukocyte antigen
HPRC: Human Pangenome Reference Consortium
HRRT: human read removal tool
MTB: *Mycobacterium tuberculosis*
MTBC: *Mycobacterium tuberculosis* Complex
NTM: non-tuberculous mycobacteria
TP: true positive
WGS: whole-genome sequencing

## Ethical Approval

Not applicable.

## Consent for publication

Not applicable

## Author’s Contributions

M.B.H.: conceptualisation, data curation, formal analysis, investigation, methodology, resources, software, writing – original draft, writing – review and editing. L.J.M.C.: funding acquisition, methodology, supervision, writing – review and editing.

## Acknowledgements

This research was supported by The University of Melbourne’s Research Computing Services and the Petascale Campus Initiative.

## References

1. Global tuberculosis report 2022. Geneva: World Health Organization; 2022.

2. Pai M, Behr MA, Dowdy D, Dheda K, Divangahi M, Boehme CC, et al. Tuberculosis. Nature Reviews Disease Primers 2016;2(1):16076. 10.1038/nrdp.2016.76.

3. Gordon AK, Marais B, Walker TM, Sintchenko V. Clinical and public health utility of Mycobacterium tuberculosis whole genome sequencing. International Journal of Infectious Diseases 2021;10.1016/j.ijid.2021.02.114.

4. Hall MB, Rabodoarivelo MS, Koch A, Dippenaar A, George S, Grobbelaar M, et al. Evaluation of Nanopore sequencing for Mycobacterium tuberculosis drug susceptibility testing and outbreak investigation: a genomic analysis. The Lancet Microbe 2022 Dec;4(e84-e92). 10.1016/S2666-5247(22)00301-9.

5. Walker TM, Lalor MK, Broda A, Ortega LS, Morgan M, Parker L, et al. Assessment of Mycobacterium tuberculosis transmission in Oxfordshire, UK, 2007–12, with whole pathogen genome sequences: an observational study. The Lancet Respiratory Medicine 2014;2(4):285–292. 10.1016/s2213-2600(14)70027-x.

6. Smith C, Halse TA, Shea J, Modestil H, Fowler RC, Musser KA, et al. Assessing Nanopore sequencing for clinical diagnostics: A comparison of NGS methods for Mycobacterium tuberculosis. Journal of Clinical Microbiology 2020;59(1). 10.1128/jcm.00583-20.

7. McNerney R, Clark TG, Campino S, Rodrigues C, Dolinger D, Smith L, et al. Removing the bottleneck in whole genome sequencing of Mycobacterium tuberculosis for rapid drug resistance analysis: a call to action. International Journal of Infectious Diseases 2017 Mar;56:130–135. 10.1016/j.ijid.2016.11.422.

8. Nilgiriwala K, Rabodoarivelo MS, Hall MB, Patel G, Mandal A, Mishra S, et al. Genomic Sequencing from Sputum for Tuberculosis Disease Diagnosis, Lineage Determination, and Drug Susceptibility Prediction. Journal of Clinical Microbiology 2023;61(3):e01578–22. 10.1128/jcm.01578-22.

9. Goig GA, Blanco S, Garcia-Basteiro AL, Comas I. Contaminant DNA in bacterial sequencing experiments is a major source of false genetic variability. BMC Biology 2020 Mar;18(1):24. 10.1186/s12915-020-0748-z.

10. Wood DE, Lu J, Langmead B. Improved metagenomic analysis with Kraken 2. Genome Biology 2019 Nov;20(1):257. 10.1186/s13059-019-1891-0.

11. Wyllie DH, Sanderson N, Myers R, Peto T, Robinson E, Crook DW, et al. Control of Artifactual Variation in Reported Intersample Relatedness during Clinical Use of a Mycobacterium tuberculosis Sequencing Pipeline. Journal of Clinical Microbiology 2018 Jul;56(8):e00104–18. 10.1128/JCM.00104-18.

12. Heupink TH, Verboven L, Warren RM, Van Rie A. Comprehensive and accurate genetic variant identification from contaminated and low-coverage Mycobacterium tuberculosis whole genome sequencing data. Microbial Genomics 2021;7(11):000689. 10.1099/mgen.0.000689.

13. Jajou R, Kohl TA, Walker T, Norman A, Cirillo DM, Tagliani E, et al. Towards standardisation: comparison of five whole genome sequencing (WGS) analysis pipelines for detection of epidemiologically linked tuberculosis cases. Eurosurveillance 2019 Dec;24(50):1900130. 10.2807/1560-7917.ES.2019.24.50.1900130.

14. The CRyPTIC Consortium and the 100,000 Genomes Project. A data compendium associating the genomes of 12,289 Mycobacterium tuberculosis isolates with quantitative resistance phenotypes to 13 antibiotics. PLOS Biology 2022 Aug;20(8):e3001721. 10.1371/journal.pbio.3001721.

15. Cuevas-Córdoba B, Fresno C, Haase-Hernández JI, Barbosa-Amezcua M, Mata-Rocha M, Muñoz-Torrico M, et al. A bioinformatics pipeline for Mycobacterium tuberculosis sequencing that cleans contaminant reads from sputum samples. PLOS ONE 2021 Oct;16(10):e0258774. 10.1371/journal.pone.0258774.

16. Mariner-Llicer C, Goig GA, Zaragoza-Infante L, Torres-Puente M, Villamayor L, Navarro D, et al. Accuracy of an amplicon-sequencing nanopore approach to identify variants in tuberculosis drug-resistance-associated genes. Microbial Genomics 2021;7(12):000740. 10.1099/mgen.0.000740.

17. Hall MB, Simulated Nanopore metagenomic reads. Zenodo; 2023. 10.5281/zenodo.8339789.

18. Hall MB, Simulated Illumina metagenomic reads. Zenodo; 2023. 10.5281/zenodo.8339791.

19. Katz KS, Shutov O, Lapoint R, Kimelman M, Brister JR, O’Sullivan C. STAT: a fast, scalable, MinHash-based k-mer tool to assess Sequence Read Archive next-generation sequence submissions. Genome Biology 2021 Sep;22(1):270. 10.1186/s13059-021-02490-0.

20. Hall MB, Kraken2 Human database. Zenodo; 2023. 10.5281/zenodo.8339700.

21. Liao WW, Asri M, Ebler J, Doerr D, Haukness M, Hickey G, et al. A draft human pangenome reference. Nature 2023 May;617(7960):312–324. 10.1038/s41586-023-05896-x.

22. Hall MB, Kraken2 Human Pangenome Reference Consortium database. Zenodo; 2023. 10.5281/zenodo.8339732.

23. Li H. Minimap2: pairwise alignment for nucleotide sequences. Bioinformatics 2018;34(18):3094–3100. 10.1093/bioinformatics/bty191, _eprint: 1708.01492.

24. Constantinides B, Hunt M, Crook DW. Hostile: accurate host decontamination of microbial sequences. bioRxiv 2023;10.1101/2023.07.04.547735.

25. Jain C, Rhie A, Hansen NF, Koren S, Phillippy AM. Long-read mapping to repetitive reference sequences using Winnowmap2. Nature Methods 2022 Jun;19(6):705–710. 10.1038/s41592-022-01457-8.

26. Rhie A, Nurk S, Cechova M, Hoyt SJ, Taylor DJ, Altemose N, et al. The complete sequence of a human Y chromosome. Nature 2023 Sep;621(7978):344–354. 10.1038/s41586-023-06457-y.

27. Kim Hs, Jeon S, Kim Y, Kim C, Bhak J, Bhak J. KOREF_S1: phased, parental trio-binned Korean reference genome using long reads and Hi-C sequencing methods. GigaScience 2022 Jan;11:giac022. 10.1093/gigascience/giac022.

28. Hall MB, Mycobacterium representative kraken2 database. Zenodo; 2023. 10.5281/zenodo.8339822.

29. Letcher B, Hunt M, Iqbal Z. Gramtools enables multiscale variation analysis with genome graphs. Genome Biology 2021;22(1):259. 10.1186/s13059-021-02474-0.

30. Hall MB, Mycobacterium tuberculosis database. Zenodo; 2023. 10.5281/zenodo.8339948.

31. Hall MB, Mycobacterium genus database. Zenodo; 2023. 10.5281/zenodo.8339941.

32. Hall MB, Clockwork database. Zenodo; 2023. 10.5281/zenodo.8339803.

33. Meehan CJ, Barco RA, Loh YHE, Cogneau S, Rigouts L. Reconstituting the genus Mycobacterium. International Journal of Systematic and Evolutionary Microbiology 2021 Sep;71(9):004922. 10.1099/ijsem.0.004922.

34. Tortoli E, Brown-Elliott BA, Chalmers JD, Cirillo DM, Daley CL, Emler S, et al. Same meat, different gravy: ignore the new names of mycobacteria. European Respiratory Journal 2019 Jul;54(1). 10.1183/13993003.00795-2019.

35. Wick R, rrwick/Assembly-Dereplicator: Assembly Dereplicator v0.3.1. Zenodo; 2023. 10.5281/zenodo.7894123.

36. Wick RR. Badread: simulation of error-prone long reads. Journal of Open Source Software 2019 Apr;4(36):1316. 10.21105/joss.01316.

37. Huang W, Li L, Myers JR, Marth GT. ART: a next-generation sequencing read simulator. Bioinformatics 2012 Feb;28(4):593–594. 10.1093/bioinformatics/btr708.

38. Langmead B, Salzberg SL. Fast gapped-read alignment with Bowtie 2. Nature Methods 2012 Apr;9(4):357–359. 10.1038/nmeth.1923.

39. Langmead B, Kraken 2 / Bracken Refseq indexes; 2023. https://genome-idx.s3.amazonaws.com/kraken/k2_standard_20230605.tar.gz.

40. Langmead B, Kraken 2 / Bracken Refseq indexes; 2023. https://genome-idx.s3.amazonaws.com/kraken/k2_standard_08gb_20230605.tar.gz.

41. Vitor C Piro, genome_updater. GitHub; 2023. https://github.com/pirovc/genome_updater.

